# Nuclear auxin signalling induces autophagy for developmental reprogramming

**DOI:** 10.1101/2025.11.04.686542

**Authors:** Caterina Giannini, Christian Löfke, Geraldine Brunoud, Enric Bertran Garcia de Ollala, Bin Guan, Stefan Riegler, Anastasia Teplova, Andres Perez Gonzalez, Marintia M. Nava García, Eva Benkova, Teva Vernoux, Yasin Dagdas, Jiří Friml

**Affiliations:** Institute of Science and Technology Austria (ISTA), 3400 Klosterneuburg, Austria; Gregor Mendel Institute of Molecular Plant Biology, 1030 Vienna, Austria; Laboratoire Reproduction et Développement des Plantes, Université de Lyon, ENS de Lyon, CNRS, INRAE, Lyon, France; Heidelberg University, Centre for Organismal Studies (COS), 69120 Heidelberg, Germany

## Abstract

The phytohormone auxin is a central regulator of plant growth and development, traditionally known for mediating transcriptional reprogramming through the canonical TIR1/AFB-Aux/IAA signalling pathway. In this study, we reveal that auxin rapidly induces macroautophagy, a catabolic process critical for the removal and recycling of superfluous macromolecules. We demonstrate that natural auxin (IAA) triggers autophagy at physiological concentrations. Genetic and pharmacological analyses show that TIR1/AFB receptors and their adenylate cyclase activity are indispensable for autophagy induction. Furthermore, auxin-induced autophagy depends on transcription, highlighting its integration with the broader gene regulatory networks. Functionally, we show that auxin-induced autophagy is required for efficient developmental reprogramming and organogenesis in both root and shoots. Autophagy is induced at places of local auxin maxima and autophagy-deficient mutants exhibit delayed differentiation and retarded organ primordia progression at the meristematic zones. Together, our findings uncover a dual role for auxin in coordinating gene expression and autophagic clearance, thereby facilitating rapid and effective developmental transitions.

## Introduction

Plants, as sessile organisms, face the challenge of adapting to a constantly changing environment without the ability to physically relocate. To overcome this limitation, they have evolved intricate regulatory systems that allow them to sense, integrate, and respond to diverse environmental and endogenous cues with remarkable plasticity.

A central component of this regulatory complexity is a class of small, mobile signalling molecules known as phytohormones^1,2^. Among these, auxin stands out as the master regulator of plant growth and development, influencing a wide array of physiological and cellular processes, from embryogenesis and organ patterning to cell division, polarization and elongation^3^.

The major role of auxin is inducing transcriptional reprogramming through the nuclear TIR1/AFB-Aux/IAA mechanism^4^. In this pathway, auxin facilitates the interaction between TIR1/AFB receptors and Aux/IAA repressors, promoting the latter’s ubiquitination and degradation via the proteasome. This degradation relieves the repression on class A auxin response factors (ARFs), and, together with TIR1/AFB-mediated production of cAMP second messenger^5^, allows for transcription of auxin-responsive genes. While this transcriptional module is essential for majority of auxin-mediated developmental responses, accumulating evidence suggests that auxin also activates rapid, non-transcriptional responses. Part of these responses are mediated by the cytoplasmic TIR1/AFB auxin receptors; chiefly by AFB1^6–8^ or by extracellular auxin signalling ABP1/ABL auxin receptors and associated transmembrane kinases such as TMK1^9–11^.

An intriguing and emerging question in auxin signalling is whether and how it crosstalks with macroautophagy^12,13^ (hereafter referred to as autophagy), a conserved catabolic process that facilitates cellular remodelling and recycling^14^. Autophagy degrades damaged organelles, misfolded proteins, and other unwanted cytoplasmic components by sequestering them at double-membraned autophagosomes and delivering them to the vacuole^14^. Autophagy has been studied predominantly in the context of stress responses, including nutrient deprivation, oxidative stress, and pathogen attack^15^. However, a vital role for autophagy in plant development starts to emerge, particularly in dynamic processes requiring rapid cellular remodelling^16,12^.

Here we demonstrate that auxin induces autophagy via the TIR1/AFB-signalling pathway and cAMP production. We show that this auxin-signalling-autophagy module is required for meristem differentiation and organogenesis in both roots and shoots. Our findings suggest that auxin not only activates new genetic programs but also triggers the active removal of obsolete cytoplasmic content, thus facilitating efficient reprogramming.

## Results

### Auxin specifically induces autophagy

High-concentration of synthetic auxin NAA, among many other stimuli, has been suggested to induce autophagy^12^. Therefore, we explored whether natural auxin IAA, at physiological concentrations, constitutes a relevant stimulus to trigger autophagy *in vivo*.

To monitor autophagy, we used the *UBI::GFP-ATG8A* line, in which the formation of puncta in the cytoplasm reports the induction of autophagy^17–19^. When we treated roots exogenously with IAA (50 nM), we observed a significant induction of puncta formation (Fig. 1a - snapshots at 1.5 h after treatment). We found the accumulation of puncta is specific to IAA, since chemically related compounds, Benzoic Acid (BA) and Tryptophan (Trp), did not induce autophagic puncta formation (Fig. 1a, Supplemental Fig. S1). We also monitored all different isoforms of ATG8 A-I and found a consistent IAA effect on puncta formation in their respective native promoter marker lines *pATG8x::GFP-ATG8x* (Supplemental videos V1-V9).

**Figure 1:**
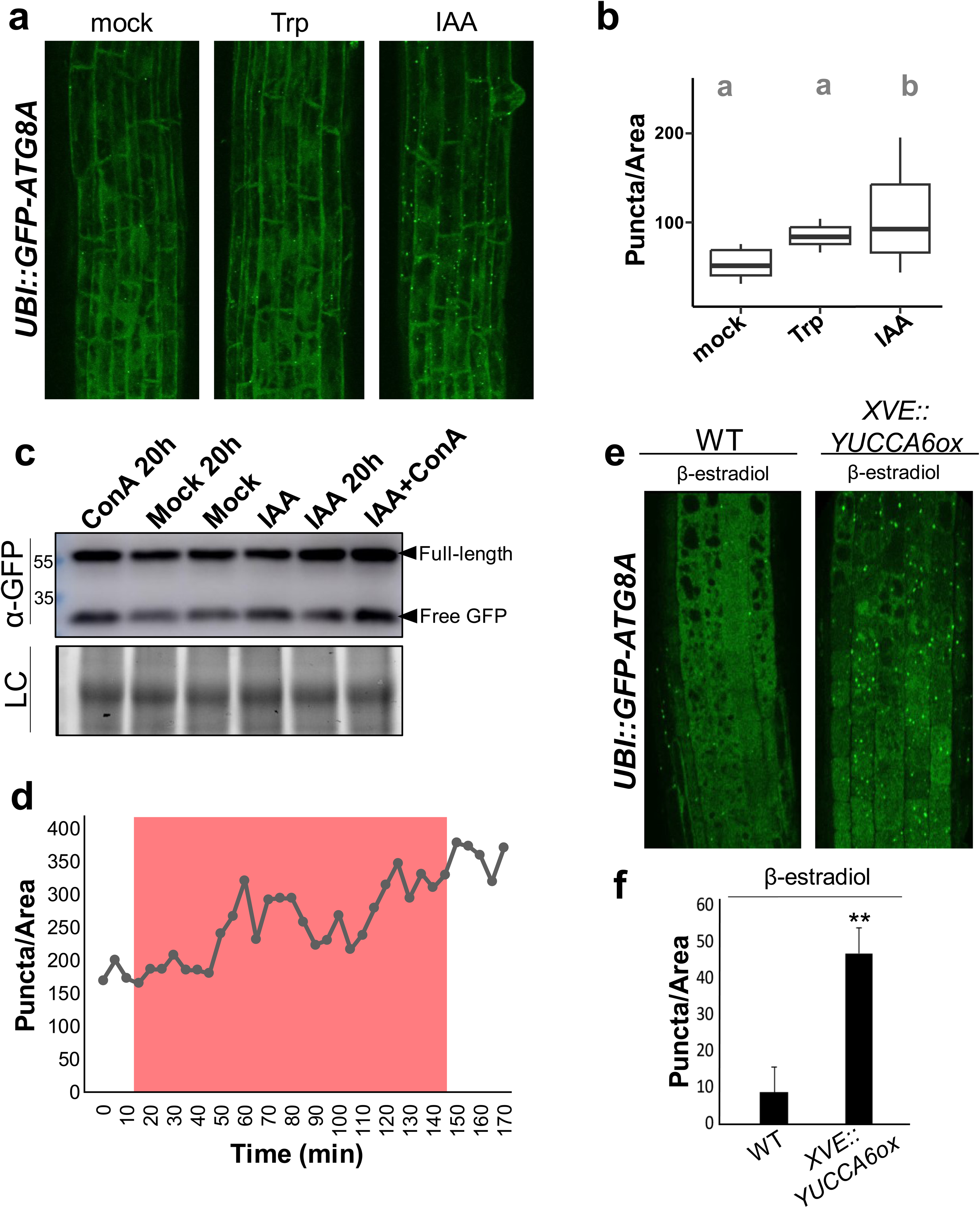
Exogenously applied or endogenously elevated auxins specifically induce autophagy. **a**, Maximal projection micrographs (1.5h) of GFP-ATG8A in roots treated with Tryptophan (1μM) and IAA (50nM) compared to mock treatment. Puncta accumulation is quantified in **b**, using Anova Post Hoc test with Turkey correction. When two genotypes share a letter, p > 0.05. **c**, Western blot based autophagic flux analysis of GFP-ATG8A samples. Free GFP band indicates induction of autophagy. 5-day old seedlings were treated with mock or indicated treatment conditions. **d**, Number of puncta over time in microfluidics set up, red window indicates IAA (50nM) treatment. Quantification of Supplemental time-lapse video V1. **e**, GFP-ATG8A maximal projection in roots of WT and *XVE::YUCCAox*, treated with β-estradiol (10μM), to induce endogenous elevated auxin level in the latter. **f**, Puncta quantification of e using Anova Post Hoc test with Turkey correction, p < 0.01.

To verify the IAA effect on autophagy by independent methods, we tested autophagic flux through western blot analysis of GFP-ATG8A^17,18^. Autophagic flux is increased following IAA treatment (50 nM for 2 h), compared to the mock conditions, as shown by increased intensity of the free GFP band (Fig. 1c). In addition, Concanamycin A (vacuolar ATPases inhibitor), which prevents degradation of the autophagic bodies in the vacuole^17^, led to an increase in both full length GFP-ATG8 and free GFP band (Fig. 1c), consistent with increased autophagic flux upon IAA treatment. These results reveal that natural auxin at physiological concentrations induces autophagy and consequent degradation at the vacuole.

To examine the dynamics of the IAA effect on autophagy, we used a microfluidics (root chip) set up^6,20^. We observed a clear induction of GFP-ATG8A puncta as early as 20-30min from the onset of the IAA treatment (Fig. 1d, Supplemental video V10).

To test the effect of endogenously produced auxin, we induced auxin biosynthesis through inducible expression of YUCCA auxin biosynthetic enzyme using *XVE::YUCCAox* transgenic line^21^. Following 4 h β-estradiol treatment, we observed a strong activation of autophagy (Fig. 1e,f).

These observations show that exogenously applied or endogenously elevated auxin induces autophagy within ∼20min.

### TIR1/AFBs auxin receptors mediate autophagy induction

Next, we asked which auxin-signalling pathway is responsible for inducing autophagy. We first used a pharmacological approach by treating with PEO-IAA, a competitive inhibitor of the TIR1 receptor^22^. When *XVE::YUCCAox* seedlings were induced to produce auxin and subsequently treated with PEO-IAA, the formation of autophagic puncta was strongly reduced. This indicates that autophagy activation depends on the canonical, TIR1-mediated auxin signalling pathway (Supplemental Fig. S1).

To confirm this result with a genetic approach, we introduced the autophagy marker *UBI::GFP-ATG8A* into the *tir triple* (*tir1afb2afb3*) mutant. We failed to observe the IAA-induced autophagy promotion in this background. Notably, autophagy induction by the salt stress (150 mM NaCl) in the *tir triple* roots occurred normally, showing that autophagy in response to auxin is impaired but that autophagy *per se* is still functional (Fig. 2a).

**Figure 2:**
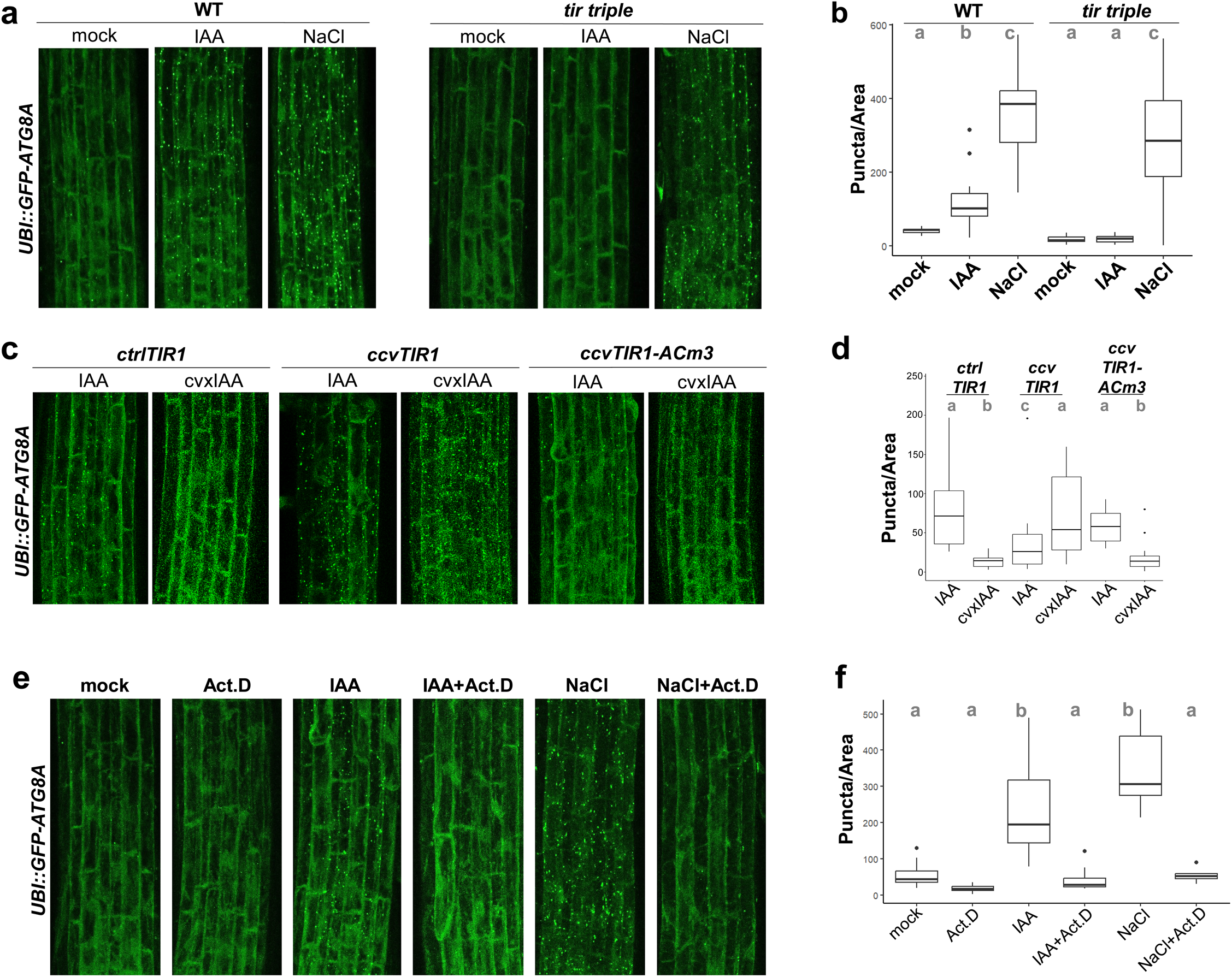
Auxin activates autophagy via the canonical TIR1/AFB-dependent signalling pathway, requiring cAMP production and transcriptional regulation. **a**, Snapshot (1.5h) of GFP-ATG8A maximal projection in WT or *tir1afb2afb3* roots treated with IAA (50nM) and NaCl (150mM) compared to WT. Puncta accumulation is quantified in **b**, using Anova Post Hoc test with Turkey correction. When two genotypes share a letter, p > 0.05. **c**, Snapshot (1.5h) of GFP-ATG8A maximal projection in F1 *crtlTIR1*, *ccvTIR1* and *ccvTIR1^ACm3^* roots treated with IAA (50nM) and cvxIAA (1μM). Puncta accumulation is quantified in **d**, using Anova Post Hoc test with Turkey correction. When two genotypes share a letter, p > 0.05. **e**, Snapshot (1.5h) of GFP-ATG8A maximal projection in roots treated with IAA (50nM), NaCl (150mM), Actinomycin D (Act. D 50 μM) and co-treatments compared to mock treatment. Puncta accumulation is quantified in **f**, using Anova Post Hoc test with Turkey correction. When two genotypes share a letter, p > 0.05.

Furthermore, we exploited the genetically engineered ccvTIR1 system that selectively activates TIR1-mediated signalling in response to the auxin analogue, cvxIAA^23^. cvxIAA treatment of ccvTIR1 plants leads to autophagy activation as manifested by increased GFP-ATG8A puncta formation (Fig. 2c). On the contrary, the *ctrlTIR1* background, which expresses the wild type (WT) receptor TIR1, as well as the *GFP-ATG8A* line in WT background, showed no autophagy activation following cvxIAA treatment (Fig. 2c, Supplemental Fig. S1c). These results confirm the specificity of TIR1-dependence in auxin-triggered autophagy. With the same approach, we tested the involvement of the dominantly cytosolic auxin receptor AFB1, which mediates rapid, non-transcriptional auxin effects on root growth^6^. The cvxIAA treatment of the ccvAFB1 line did not lead to any GFP-ATG8A accumulation, suggesting AFB1 is not involved in activating autophagy in response to auxin (Supplemental Fig. S2a,b).

In addition, we also tested GFP-ATG8A in *tmk1-1*, *abp1-C1* and *abp1-TD1* backgrounds to assess any contribution of auxin cell surface signalling pathway for auxin-triggered autophagy. The results show that auxin induces autophagy in these mutants as well as in WT, arguing against the dependence to the cell surface signalling (Fig. S2c,d).

Altogether, these pharmacological and genetic approaches revealed that auxin induces autophagy via the nuclear TIR1/AFB-based signalling pathway.

### Transcriptional mechanism downstream of TIR1 adenylate cyclase activity induces autophagy

Recent studies showed that TIR1/AFBs receptors have adenylate cyclase (AC) activity that is required for transcriptional regulation in response to auxin^24,25^. We tested whether the production of cAMP by TIR1 is required for the autophagy induction by auxin. While the ccvTIR1 showed normal autophagy activation after treatment with cvxIAA, in the ccvTIR1^ACm3^ line with compromised AC activity^25^, the auxin-induced autophagy activation was absent (Fig. 2c, Supplemental Fig. S1) showing that TIR1-mediated cAMP production is indispensable for autophagy induction.

Specific involvement of TIR1 and its AC activity thus suggests that auxin regulates autophagy by a transcriptional mechanism. We further tested this idea using Actinomycin D (Act D) and cycloheximide (CHX) treatments. Act D inhibits transcription, whereas CHX blocks translation. We observed that Act D and CHX alone do not activate autophagy, but when co-applied with IAA, they block accumulation of autophagic puncta (Fig. 2e, Supplemental Fig. S1e). Moreover, we found that Act D and CHX block autophagy even in salt stress conditions (150 mM NaCl). This suggests that autophagy activation in general depends on transcription, not only in the context of auxin but also in other instances.

These observations show that autophagy is induced by a transcription-dependent mechanism, which in the case of auxin is downstream of TIR1/AFB-mediated cAMP production that is required to couple auxin transcriptional responses to an activation of autophagy.

### Autophagy correlates with and is required for root development and organogenesis

Our findings show that TIR1/AFB-mediated auxin signalling does not only mediate transcriptional reprogramming^4,25^ but also simultaneously induce autophagy. Therefore, we tested whether this autophagy induction plays a role in auxin-induced developmental transitions using the recently discovered autophagy receptor CESAR2-GFP^26^ and autophagy-deficient mutants *atg2-1* and *atg5*-*1*^27,28^.

First, we looked at progression of differentiation in root apical meristem, a well-established model of continuous cell division and differentiation. Auxin gradients within the root meristem are known to determine whether cells remain in a meristematic state or begin to specialize into different root cell types^29,30^. To test whether auxin-regulated autophagy may contribute to this transition, we analysed meristem architecture in *atg2* and *atg5* mutants. Measuring the length of the root meristem zone, we found that autophagy mutants have an extended meristem length, suggesting more cells do not undergoing transition to differentiation compared to WT (Fig. 3a). This indicates that developmental transition to differentiated cells in root apical meristem is delayed when autophagy is compromised.

**Figure 3:**
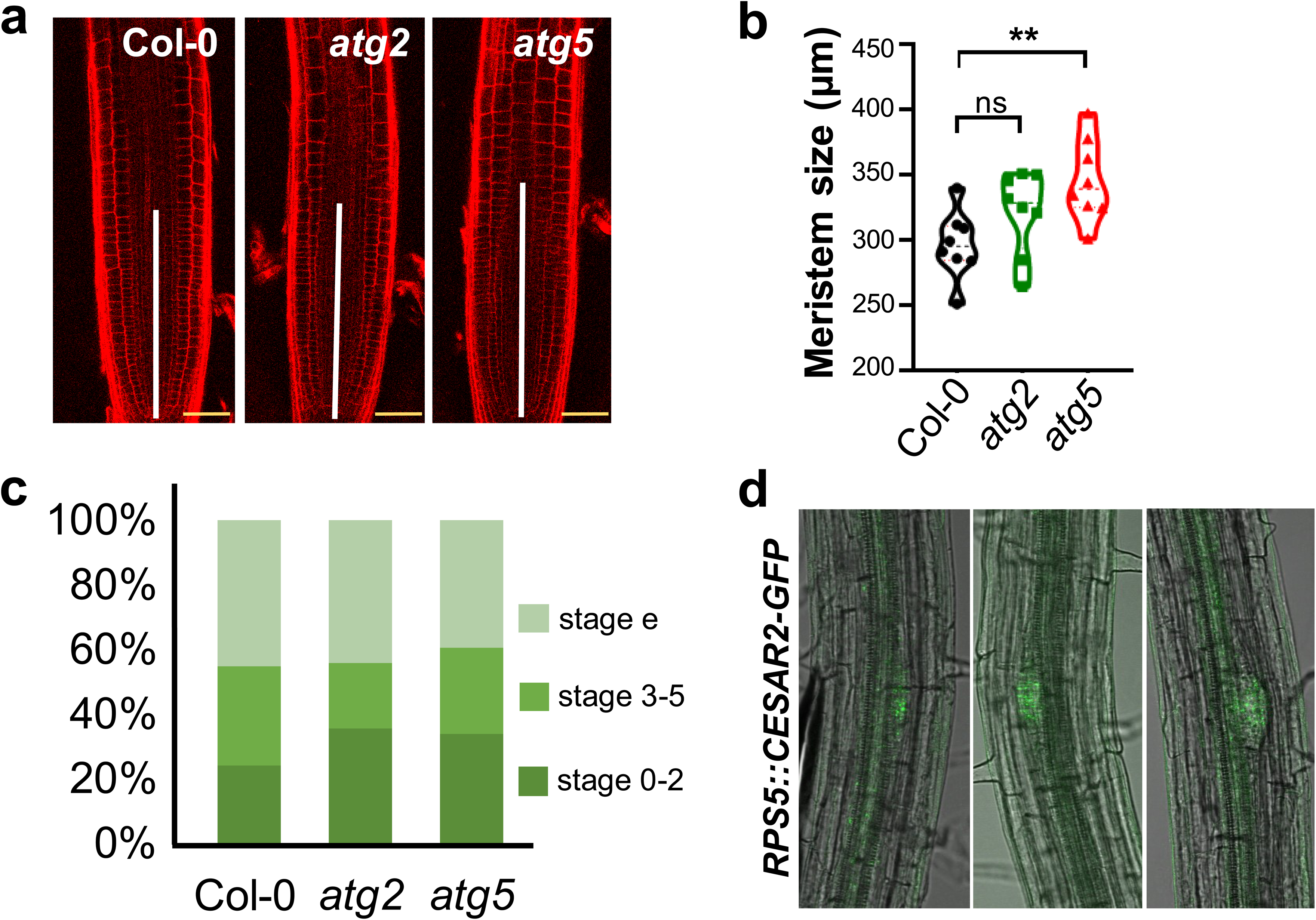
Autophagy is required for auxin-regulated developmental reprogramming in root meristem differentiation and lateral root organogenesis. **a**, PI stained roots of WT, *atg2* and *atg5* mutants, highlighting their meristem size (white line). Scale bar in yellow is 50μm. **b**, Quantification of meristem length, comparisons using Anova Post Hoc test with Turkey correction p < 0,001. **c**, Lateral root primordia progression analysis for WT, *atg2* and *atg5* mutants. WT=11, *atg2*=10, *atg5*=9. **d**, Expression pattern of translational fusion of CESAR2-GFP marker (n=15), maximal projection of GFP signal and bright field (BF) overlay of 6-7 days old seedling roots. CESAR2-GFP puncta accumulates in founder cells and lateral root primordia. Time-lapse of CESAR2-GFP signal from founder cell to lateral root primordia progression is shown in Supplemental video V2.

Lateral root formation is another classical auxin-induced process dependent on TIR1/AFB1 signalling and involving extensive developmental reprograming. It has been shown that during lateral root development auxin accumulates already in primordia founder cells and later the auxin maximum at the primordia tip guides lateral root primordia progression^31^. Accordingly, overnight time-lapse imaging of CESAR2-GFP revealed puncta accumulation in lateral root primordia and distinct pericycle cells, which were identified as lateral root founder cells by live imaging, as they eventually formed lateral root primordia (Fig. 3d - different LR primordia from different roots, Supplemental video V11 of CESAR-GFP time-lapse during LR primordia development).

Consistently, a previous study showed that *atg2-1* and *atg5-1* mutants have lower lateral root (LR) density^13^. We examined in detail whether autophagy is required for lateral root organogenesis by analysing its progression in *atg2* and *atg5* mutants. The autophagy-deficient mutants showed a similar number of primordia overall and a comparable primary root length as the WT seedlings. However, *atg2* and *atg5* seedlings showed a clear trend of delayed developmental progression of the primordia as compared to WT. In particular, many primordia were retained at early stages (stage 0-2, 33-34% vs 23% of WT), consistent with delayed developmental transitioning during lateral root formation.

These results suggest that autophagy is required for auxin-induced developmental reprogramming during root meristem differentiation and lateral root primordia development.

### Autophagy correlates with and is required for shoot-derived organogenesis

Next, we tested whether auxin-regulated autophagy played a more general role in plant development by analysing its contribution to shoot meristem differentiation and shoot-derived organogenesis.

Imaging of CESAR2-GFP in the shoot apical meristem showed puncta accumulation in regions of endogenous high auxin accumulation, correlating with sites of new organ formation^3,31^ (Fig. 4a,b). In addition, we grew CESAR2-GFP plants on the auxin efflux inhibitor NPA^32^ to prevent organogenesis and analysed the recovery of organogenesis after the release of shoot apical meristem from the NPA^33^. In these conditions, CESAR2-GFP accumulated in puncta in the areas with a dynamics similar to a marker for transcriptional auxin signalling (DR5) that precedes new organogenesis, and again marked new lateral organs being produced (Fig. 4c). These observations suggest that also in the context of the shoot-derived organogenesis, the cells, where auxin is known to accumulate, show increased autophagy rates.

**Figure 4:**
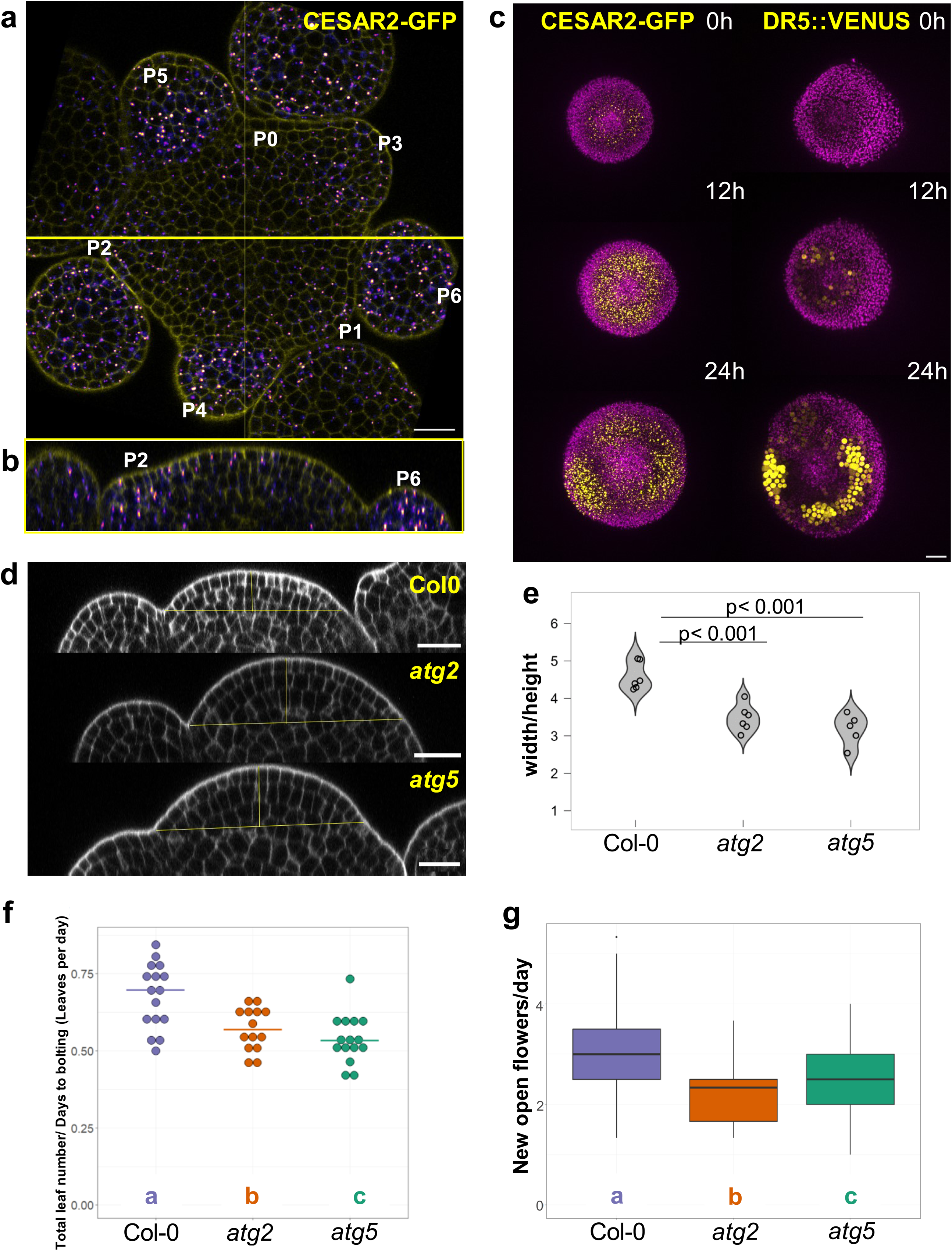
Autophagy contributes to lateral organ formation in shoots. Expression pattern of CESAR2-GFP in SAM (n=16) **a**, maximal projection **b**,orthogonal view following the yellow line on maximal projection, cell walls stained by PI are in yellow, CESAR-GFP fusion proteins are in LUT Fire. **c**, Time course of organogenesis recovery in SAM after NPA treatment, translational fusion of CESAR2-GFP marker (n=12) and DR5::3xVenusN7 (n=4), signal Venus and GFP are in yellow and autofluorescence is in magenta. **d**, Side views of inflorescence meristem of WT (n=6), *atg2* (n=6), and *atg5* (n=6) mutants, cell wall stained by PI are in grey, **e**, quantification of width/height ratio, comparisons using Anova Post Hoc test with Turkey correction p < 0,001. **f**, Plastochron during vegetative development, n=15 plants, comparisons using Wilcoxon rank-sum test with Benjamini-Hochberg corrections, when two genotypes share a letter, p > 0.05 and **g**, reproductive development, number of calculated rates (WT =131, *atg2* = 74, *atg5* = 86), comparisons using Wilcoxon rank-sum test with Benjamini-Hochberg corrections, when two genotypes share a letter, p > 0.05. Scale bar for all images: 20µm.

Analysis of the shoot apical meristem in *atg2* and *atg5* plants revealed a change in the shape of the meristem, the mutant meristems being more domed, notably to an increase in height (Fig. 4d, Supplemental Fig. S4). This phenotype suggests a possible perturbation of differentiation coordination at the shoot apical meristem, similar to root apical meristem. Next, we analysed shoot-derived organogenesis. Indeed, while the organs were initiated at the right positions, in *atg2* and *atg5* mutant plants, the number of leaves per days after bolting and new open flowers per day compared to WT plants was significantly decreased (Fig. 4f,g). This shows that the plastochron is slowed down, consistent with a slower dynamics of organ differentiation in these autophagy-deficient mutants.

Taken together, our observations indicate that in the shoot apex, autophagy is activated in regions marked by high auxin levels and that this activation is required for timely progression of differentiation during leaf and flower organogenesis.

## Discussion

The phytohormone auxin is the main endogenous signal mediating developmental reprograming during plant embryogenesis, organogenesis and many other developmental transitions^3,34^. It has been well established that this happens by transcriptional reprogramming mediated by the nuclear TIR1/AFB signalling^3^. This works demonstrates that this auxin signalling mechanism does not only trigger transcription of thousands of genes^35^ but simultaneously activates autophagy. Auxin thus functions not only to specify new transcriptional identities but also to trigger the clearance of pre-existing, potentially incompatible components. This dual action ensures efficient developmental reprogramming.

While this study establishes that it is the TIR1/AFB-mediated, cAMP-dependent transcriptional branch of auxin signalling that activates autophagy and demonstrates the importance of auxin-induced autophagy in root- and shoot-derived differentiation and organogenesis, the downstream mechanistic details remain unresolved. Notably, the identity of the autophagy regulators that are targeted by this mechanism remains to be established. Also, whether canonical auxin response factors (ARFs) alone mediate this induction, or whether distinct or auxiliary transcription factors are involved, is not yet clear. Additionally, it remains to be determined whether autophagy acts broadly or selectively during these developmental transitions - removing general cytoplasmic content, or targeting specific fate-incompatible regulators, protein complexes, or organelles. These questions define an important frontier in understanding how cellular identity is reprogrammed in plants.

This study also reveals that autophagy activation not only in response to auxin but also to abiotic cues - such as salt stress - requires a transcription-dependent mechanism. This suggests that autophagy induction in plants is broadly transcriptionally regulated and this might provide a general mechanism to couple transcription to developmental and stress responses. This layer of transcriptional control may have been overlooked in prior studies focused on stress or starvation responses, because autophagy is typically interpreted as a downstream metabolic adaptation. However, the reliance on transcription we identified argues for a more proactive role: autophagy is not merely a clean-up mechanism activated during crisis, but a regulated developmental tool, that can be recruited in a specific, signal-dependent manner.

Taken together, these findings propose that auxin-induced autophagy is part of a larger, conserved transcriptional module for autophagy regulation in plants. This module is engaged by both hormonal and environmental signals and may be coordinated through yet-unidentified master regulators. Clarifying the identity of these regulators and their interaction with core auxin and other signalling components will be crucial to understand how different signals induce autophagy. In parallel, uncovering the cargo specificity of autophagy during reprogramming - whether it targets repressors, organelles, or fate-inhibiting proteins - will help determine whether autophagy acts permissively or instructively during cellular transitions.

In summary, this work represents a significant conceptual advance of our understanding of how auxin mediates developmental transitions. It also has broader implications with the proposed emerging model, where autophagy is transcriptionally induced and integrated downstream of various signalling pathways, ensuring that reprogramming is both genetically instructed and cytoplasmically enabled.

## Acknowledgments

We gratefully acknowledge the support of the LSF and IOF Facilities at the Institute of Science and Technology Austria for their invaluable contributions. This project is supported by the European Research Council (ERC) under the European Union’s Horizon 2020 research and innovation program (101142681 CYNIPS) and Austrian Science Fund (FWF; P 37051-B), both to J.F.; E.B.G. and T.V. were supported by the European Union, ERC TEMPO Project 101095380 to T.V..

## Author contributions

Conceptualization, J.F., Y.D. and C.G.; initiated the project, C.L.; plant material preparation, root microscopy, LR phenotyping and imaging, image segmentation and data analysis, C.G.; SAM phenotyping and imaging, G.B.; plastochron experiments, E.B.G.; autophagic flux, B.G.; quantification software, S.R.; microfluidics time-lapse, A.T.; RAM phenotyping, C.G. and A.P.G.; autophagy methodology assistance, M.M.N.G; writing, C.G. and J.F.; reviewing and editing, all authors; and funding acquisition, J.F., T.V., E.B. and Y.D..

## Materials and Methods

### Plant materials and growth conditions

All *Arabidopsis* lines used in this study are in Col-0 background. For root experiments, all *Arabidopsis* seeds were surface-sterilized with 70% (v/v) ethanol for 10 min, followed by 100% ethanol for 10 min. Seed stratification was conducted in the dark at 4°C for 1-2 days. Seedlings were grown at 22°C on half-MS plates with 1% agar and 1% sucrose, or in soil with 16h light/8h dark cycles photoperiod at 80 to 100 mE/m^2^/sec^1^.

For shoot experiments, all *Arabidopsis* seeds were grown on soil (except for NPA experiment), after a 2-day stratification, under long-day conditions (16 h light at 22°C / 8 h dark at 20°C). For NPA experiment, seeds were surface-sterilized by gas (100 ml of bleach + 4 ml HCl 37%) for 3 h. After sterilization seeds were stratified for 2 days, the seedlings were grown on Arabidopsis medium (Duchefa, Ref: DU0742.0025) supplemented with 10 mM NPA N-(1-Naphthyl)phthalamic Acid under long-day conditions (16 h light at 22°C / 8 h dark at 20°C).

*UBI::GFP-ATG8A* line was previously reported here^19^. This marker line was crossed in different backgrounds: *tir1afb2afb3*^36^, *abp1-C1*^37^, *abp1-TD1*^37^, *tmk1-1*^38^, *ccvTIR1*^23^, *ccvTIR1^ACm1^* ^25^, *ccvTIR1^ACm3^* ^25^, *ccvAFB1*^39^, and *XVE::YUCCAox*^21^. *ATG8x:GFP-ATG8x*, *RPS5::CESAR2-GFP*, *atg2* and *atg5* lines were described respectively in^40,26–28^. *DR5::3xVenus-N7* line is the one generated in^41^.

### Plant chemical treatments for confocal microscopy

5-day-old (if not stated differently) seedlings were incubated for 1.5-2 h on half-MS 1% agar medium supplemented with different treatments: 50 nM IAA (Sigma-Aldrich), 10 μM PEO-IAA (ChemIm Ltd), 10 μM β-estradiol (Sigma-Aldrich), 1 μM L-Tryptophan (Sigma-Aldrich), 1 μM Benzoic Acid (Sigma-Aldrich), 1 μM cvxIAA (TCI, Tokyo Chemical Industry Co.), 50 μM Actinomycin D (Sigma-Aldrich), 1 μM Cycloheximide (Sigma-Aldrich), 150 mM NaCl.

### Root microscopy and imaging acquisition

Confocal microscopy was performed on a vertically mounted Zeiss LSM800 microscope equipped with an air Plan Apochromat 20 x/0.8 M27 objective. GFP-tagged proteins were excited at 488 with emission collected at range 490-576 nm. All *GFP-ATG8* lines were imaged by taking 10 Z-sections spanning 10 μM volume of each root. These were processed through “maximum intensity” projection in Fiji and puncta were quantified as descripted below (in Image segmentation for autophagic puncta quantification). For roots stained with propidium iodide (PI), fluorescence signal was detected at excitation 536 nm and emission 617 nm. Meristem size was measured manually in Fiji using the line function starting at the columella until the first cell having a width/height ratio greater than 1.

### Microfluidics setup for root time-lapse imaging

For the real-time monitoring of autophagosome formation, the Elveflow microfluidics setup coupled with a vertically mounted Zeiss LSM800 microscope was used. The microfluidics setup replicates the one described in^6^. Briefly, the seedlings are placed in the channels of the PDMS silicone chip, covered with a coverslip, and placed in the microscope with a custom-made holder. Piezo electric pressure controller (OB1 MK3+, Elveflow) coupled with flow sensors (MFS D 2+, Elveflow) and the ESI software (v.3.06.00) are used to control the flow rate and switch between control and treatment solutions. The microfluidics setup enabled the rapid exchange between the IAA treatment and the half-MS medium supplemented with an equal amount of ethanol as a control solution. 4-day-old *UBI::GFP-ATG8A* seedlings were placed on the microfluidics chip (2 seedlings per channel). It should be mentioned that during the assembly of the microfluidics chip, the seedlings are unavoidably subjected to the treatment solution mixed with the control solution. After mounting the microfluidics device and locating the holder vertically in the microscope, the seedlings were left for 60-90 min under the constant flow 5 μl/min of the control half-MS solution to recover. Then the seedlings were treated for 2 h with 50 nM IAA (constant flow 5μl/min), followed by 2 h wash out with control half-MS solution (flow 5 μl/min). Treatment/washout cycles were repeated several times. To monitor the arrival of the treatment, TRITC-Dextran dye (Sigma) (∼1 μg/ml) was added to the IAA solution. Images were taken every 5 min using air Plan Apochromat 20 x/0.8 M27 objective, 488 nm excitation laser for GFP imaging, and 561 nm excitation laser for the TRITC-dextran imaging. Z-stack spanning 20 μm of the root epidermis in the elongation zone was imaged, and for the analysis, the maximum Z-projection was used. At least 2 roots were imaged in 1 replica; 4 replicas were performed.

### Image segmentation for autophagic puncta quantification

A Tensorflow implementation of U-Net^42^, one of the most common models used for semantic segmentation tasks in biological images, was adapted from https://github.com/bnsreenu/python_for_microscopists/blob/master/204-207simple_unet_model.py. Autophagy bodies in a subset of images were manually annotated in Fiji to create binary masks for model training. Employing Tensorflow (2.17.1) and Keras (3.5.0) the model was trained on image tiles of 256×256 pixels in size, using the Adam optimizer^43^, a learning rate of 0.001, binary cross entropy as loss function, and a batch size of 16 for 50 epochs. Training as well as inference were performed in the Google Collaboratory environment. The model parameters and code for the application of the model are available upon request.

### Plant protein extraction and autophagic flux testing

For total protein extraction, *Arabidopsis* 7-day-old seedlings were harvested after treatment and ground to a fine powder in liquid nitrogen using a pre-chilled mortar and pestle. The powdered tissue was homogenized in ice-cold lysis buffer (50 mM Tris-HCl, pH 7.4, 150 mM NaCl, 1% Triton X-100, 1% sodium deoxycholate, 0.1% SDS) supplemented with protease inhibitor cocktail (Roche) and 1 mM phenylmethylsulfonyl fluoride (PMSF). The homogenate was incubated on ice for 30 min, followed by centrifugation at 12,000 × g for 30 min at 4 ℃. The clarified supernatant was transferred to a fresh microcentrifuge tube. For autophagic flux western blot analysis, total protein extracts were mixed with 4× SDS-PAGE loading buffer, boiled at 95°C for 5 min, and separated by SDS-PAGE on 4–15% gradient gels. Proteins were then electrophoretically transferred onto a polyvinylidene difluoride membrane using a wet transfer system. After transfer, membranes were blocked in 5% non-fat milk in TBS-T buffer (20 mM Tris-HCl, pH 7.5, 150 mM NaCl, 0.1% Tween-20) for 1 h at room temperature. The membranes were then incubated overnight at 4°C with the primary rabbit anti-GFP antibody (Abcam, ab290; 1:1000) diluted in the blocking buffer. After three 10 min washes with TBS-T buffer, membranes were incubated with horseradish peroxidase (HRP)-conjugated secondary antibodies (Cytiva NA9340) for 1 h at room temperature. Following 3 additional 10 min washes, the signal was detected using enhanced chemiluminescence (ECL) substrate and visualized using a digital imaging system (GE Healthcare Amersham 600).

### Lateral root primordia progression scoring

8-day-old seedlings were cleared according to the protocol as previously published^44^. Briefly, seedlings were incubated in 70% EtOH for 2 days. Afterwards, they were incubated with 4% HCl, 20% methanol at 65°C for 10 min, in 7% (w/v) NaOH, 60% ethanol at room temperature (RT) for 15 min, followed by incubation in a series of ethanol dilutions from 60% to 10% at RT. Seedlings were mounted on glass slides in drop of mounting medium (50% glycerol). The developmental stages of lateral roots (LRs) and lateral root primordia (LRP) in circa 10 seedlings per genotype were scored using Olympus BX53 microscope.

### Shoot microscopy and image acquisition

Inflorescence meristems were dissected from the main stem when its length was between 0.5 and 1.5 cm and transferred *in vitro* onto half-MS medium supplemented with vitamins and cytokinin to allow continuous growth and organogenesis, as described previously by^45^. When required, meristems were stained with 100 µM propidium iodide (Sigma Aldrich, Ref: P4864) for 5 min. Imaging was done on an inverted Zeiss LSM710 using x40-NA1 long-distance water objective with image size 1024 x 1024, pinhole at 1 AU. The laser settings were kept the same for a given reporter to allow for comparisons between samples^46^.

All image analyses were performed in Fiji. For measurement of width and height of inflorescence meristems, the position of the fourth visible primordium was used to extract an orthogonal section cutting through the centre of the meristem. The position where width and height were measured on the orthogonal section is shown in Supplemental Figure S4.

### Plastochron measurement and analysis

For vegetative plastochron measurements, the number of rosette leaves and the number of cauline leaves on the primary inflorescence was counted. Plant age was calculated from the first day the plants were exposed to light and used to score the bolting time, defined as the time point when the inflorescence had elongated 0.5 cm from the rosette. The vegetative plastochron was estimated by dividing the total leaf number by the days to bolting.

To estimate the reproductive plastochron, the new open flower rate was calculated. For this, the cumulative open flower number was recorded for each plant until proliferative arrest. At each time point, the new flower opening rate was calculated by dividing the number of new opened flowers by the number of days since the last measurement. Rates that were greater than 0 were used to compare genotypes. As measurements were not performed every day, we deleted the first and last calculated rate. The selected time window corresponds to the phase of linear growth of the cumulative open flower number.

---

**Figure S1:**
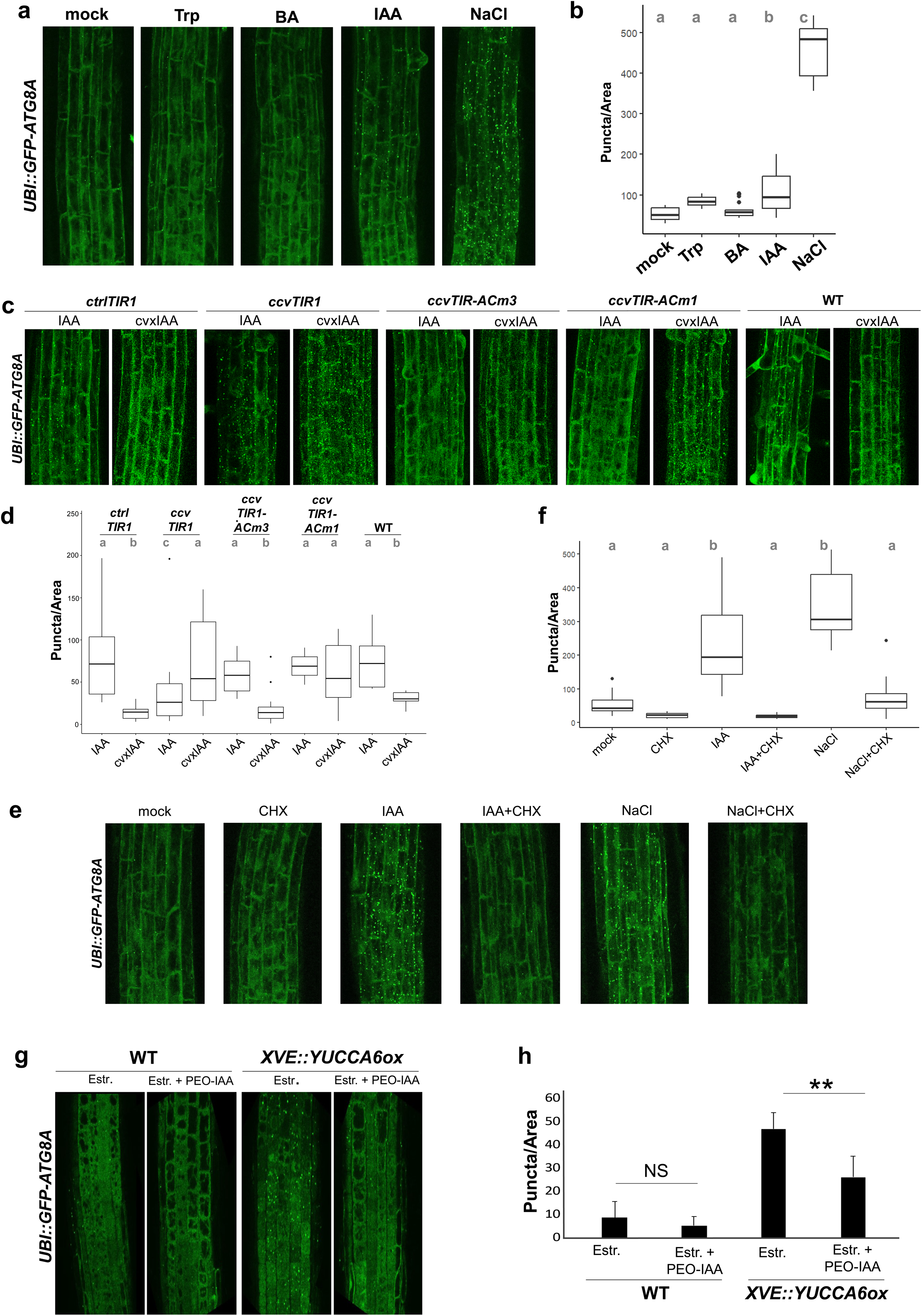
Auxin specifically induces autophagy through TIR1/AFBs pathway-extended imaging. **a**, Snapshot (1.5h) of GFP-ATG8A maximal projection in roots treated with Tryptophan (1μM), Benzoic acid (BA 1μM) and IAA (50nM) compared to WT. Puncta accumulation is quantified in **b**, using Anova Post Hoc test with Turkey correction. When two genotypes share a letter, p > 0.05. **c**, Snapshot (1.5h) of GFP-ATG8A maximal projection in F1 *crtlTIR1*, *ccvTIR1*, *ccvTIR1^ACm^*^1^ and *ccvTIR1^ACm3^* roots treated with IAA (50nM) and cvxIAA (1μM). Puncta accumulation is quantified in **d**, using Anova Post Hoc test with Turkey correction. When two genotypes share a letter, p > 0.05. **e**, Snapshot (1.5h) of GFP-ATG8A maximal projection in roots treated with Cycloheximide (CHX 1μM), IAA (50nM), NaCl (150mM) and co-treatments compared to mock treatment. Puncta accumulation is quantified in **f**, using Anova Post Hoc test with Turkey correction. When two genotypes share a letter, p > 0.05.**g**, GFP-ATG8A maximal projection in roots of WT and *XVE::YUCCAox*, treated with β-estradiol (10μM) to induce endogenous elevated auxin level in the latter and PEO-IAA (10μM) to block TIR1 pathway. **h**, Puncta quantification of g, using Anova Post Hoc test with Turkey correction, p < 0.01.

**Figure S2:**
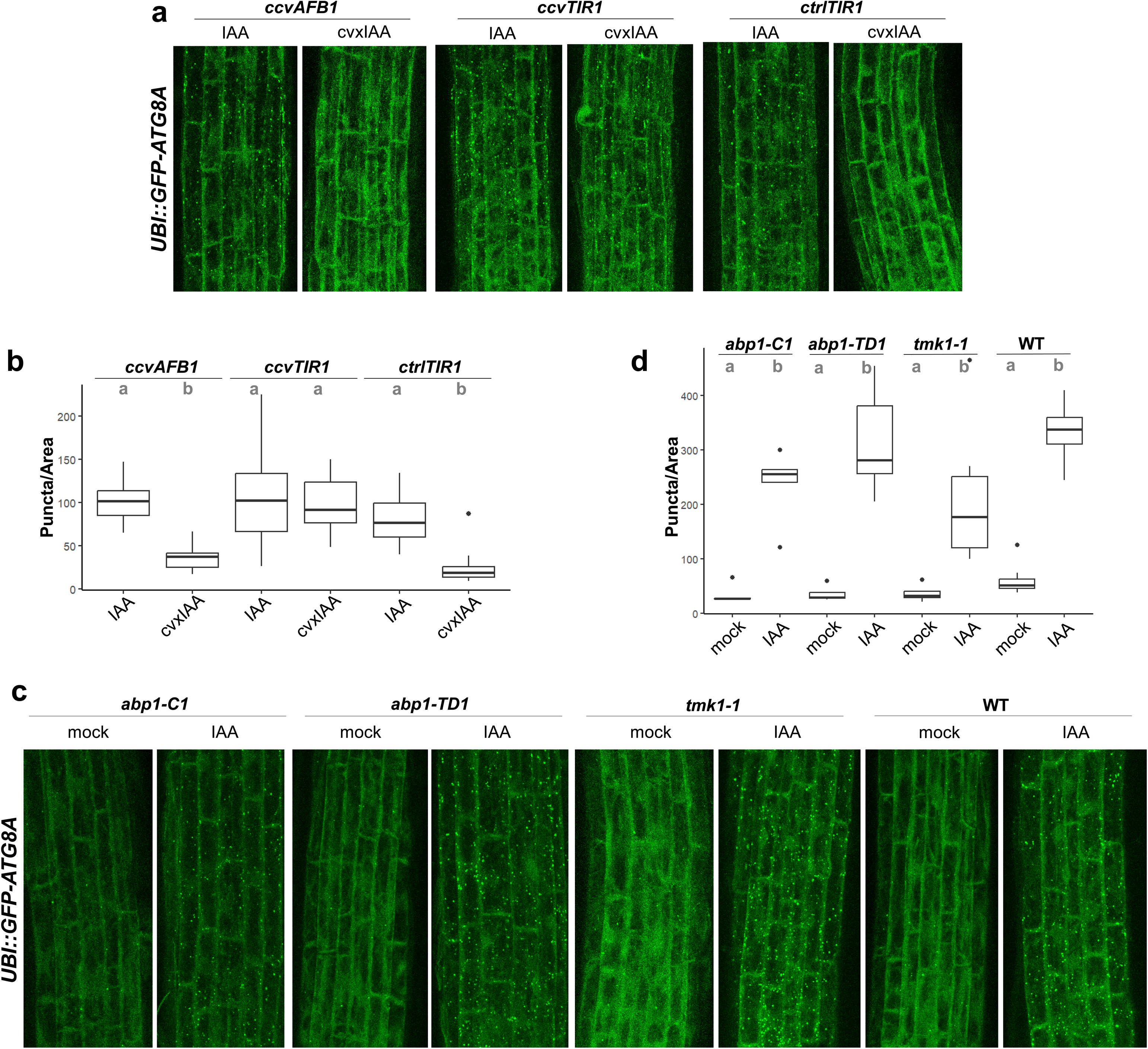
Auxin-induced autophagy does not depend on AFB1 or ABP1-TMK1 signalling pathway. **a**, Snapshot (1.5h) of GFP-ATG8A maximal projection in F1 *ccvAFB1*, *ccvTIR1* and *crtlTIR1* roots treated with IAA (50nM) and cvxIAA (1μM). Puncta accumulation is quantified in **b**, using Anova Post Hoc test with Turkey correction. When two genotypes share a letter, p > 0.05. **c**, Snapshot (1.5h) of GFP-ATG8A maximal projection in roots treated with IAA (50nM) compared to WT. Puncta accumulation is quantified in **d**, using Anova Post Hoc test with Turkey correction. When two genotypes share a letter, p > 0.05.

**Figure S3:**
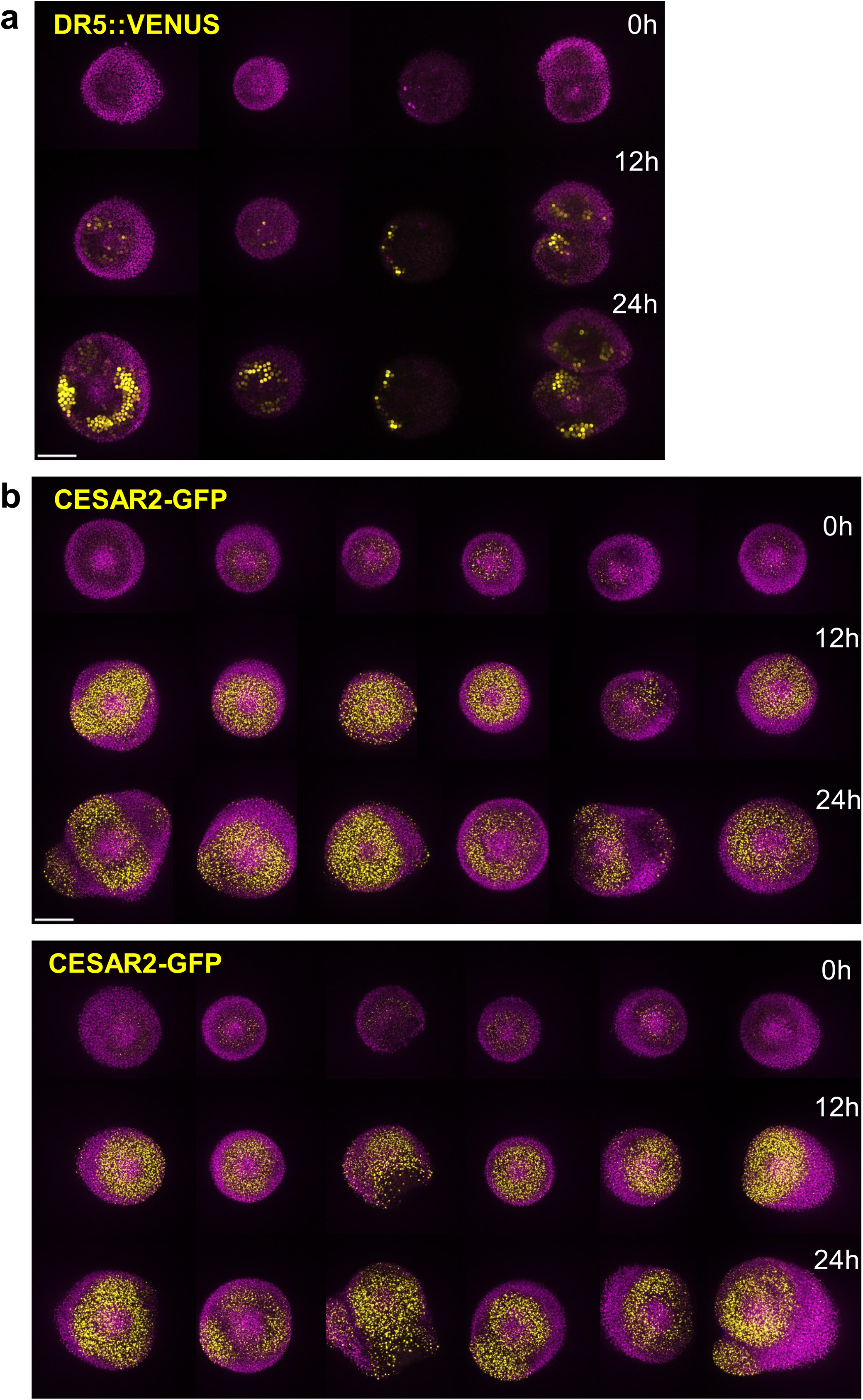
Supplemental imaging for autophagy in SAM. Time course of organogenesis recovery after NPA treatment. **a**, 4 Meristems presented in column *DR5::3xVENUS-N7* control, **b**, 12 Meristems in 2 panels *pCESAR2::CESAR-GFP*. Venus and GFP are in yellow, autofluorescence is in magenta. Scale bar is the same for all images: 50µm.

**Figure S4:**
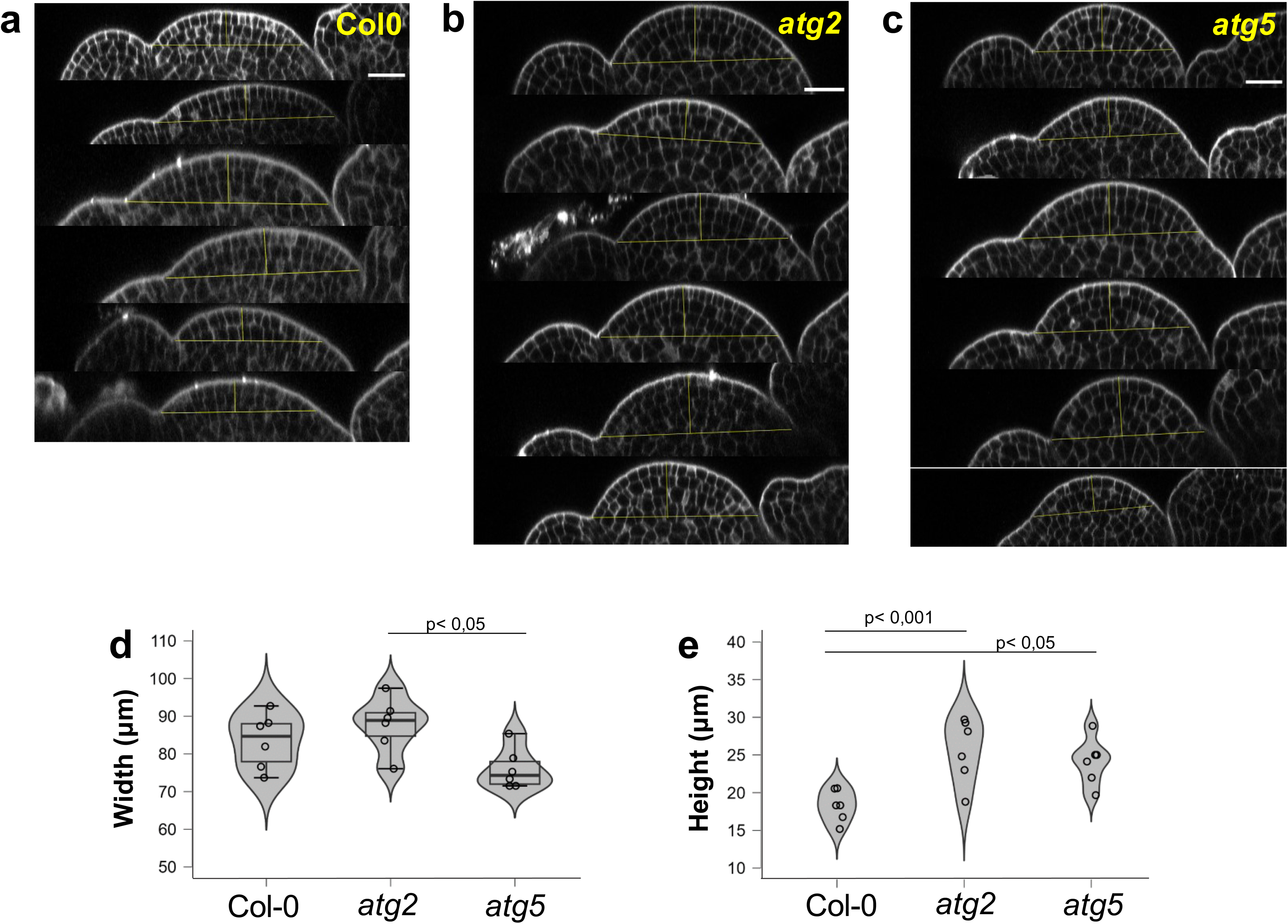
Supplemental imaging and quantification of *atg* mutants SAM. Side views of inflorescence meristem of **a**, WT (n=6), **b**, *atg2* (n=6) and **c**, *atg5* (n=6) mutants, cell wall stained by PI are in grey. Vertical and horizontal yellow bars indicate where height and width were measured, respectively. **d**, Width quantification: comparisons using Anova Post Hoc test with Turkey correction. **e**, Height quantification: comparisons using Anova Post Hoc test with Turkey correction. Scale bar is the same for all images: 20µm.

